# The Metabotropic Glutamate Receptor 5 Negative Allosteric Modulator Fenobam: Pharmacokinetics, Side Effects, and Analgesic Effects in Healthy Human Subjects

**DOI:** 10.1101/391383

**Authors:** Laura F. Cavallone, Michael C. Montana, Karen Frey, Dorina Kallogjeri, James M. Wages, Thomas. Rodebaugh, Tina Doshi, Evan D. Kharasch, Robert W. Gereau

## Abstract

Metabotropic glutamate receptor 5 (mGlu5) has been shown to modulate nociception in animals, but no mGlu5 antagonists have been developed commercially as analgesics. The mGlu5 antagonist fenobam [N-(3-chlorophenyl)-N’-(4,5-dihydro-1-methyl-4-oxo-1H-imidazole-2-yl)urea] was originally evaluated for development as a non-benzodiazepine anxiolytic. Fenobam is analgesic in numerous mouse pain models, acting exclusively via mGlu5 blockade. Furthermore, fenobam showed no signs of analgesic tolerance with up to two weeks of daily dosing in mice. Analgesic effects of fenobam in humans have not been reported. The purpose of this investigation was to evaluate fenobam pharmacokinetics and analgesic effects in humans. We first evaluated single-dose oral fenobam disposition in a parallel-group dose-escalation study in healthy volunteers. A second investigation tested the analgesic effects of fenobam in an established experimental human pain model of cutaneous sensitization utilizing capsaicin cream and heat, in a double-blind placebo-controlled study. The primary outcome measure was the area of hyperalgesia and allodynia around the area applied with heat/capsaicin. Secondary outcome measures included nociception, measured as pain rating on a visual analog scale, heat-pain detection threshold, and effects on cognition and mood. Fenobam plasma exposures showed considerable inter-individual variability, and were not linear with dose. Fenobam reduced sensitization vs placebo at a single time-point (peak plasma concentration); we found no other difference between fenobam and placebo. Our results suggest highly variable fenobam disposition, and minimal analgesic effects at the dose tested. We suggest that future studies testing analgesic effects of mGlu5 blockade are warranted, but such studies should employ molecules with improved pharmacokinetic profiles.

## Introduction

Metabotropic glutamate receptor 5 (mGlu5) has emerged as a potential candidate for the development of a new class of analgesic drugs. Despite demonstrated anti-nociceptive effects of mGlu5 antagonists in a broad range of animal pain models [5; 14], so far, no mGlu5 antagonists have been developed commercially as analgesics.

The investigational drug fenobam [N-(3-chlorophenyl)-N’-(4,5-dihydro-1-methyl-4-oxo-1H-imidazole-2-yl)urea] was originally evaluated for development by Ortho-McNeil (McN-3377) in the late 1970s as a non-benzodiazepine anxiolytic, with a then unknown molecular target[12; 15; 22; 23]. While its commercial development was not pursued, in 2005, Porter et al. characterized fenobam as a selective, non-competitive mGlu5 antagonist[25]. In agreement with earlier reports of the role of mGlu5 in nociceptive pathways[29] and the selectivity of fenobam for mGlu5, the analgesic effect of fenobam has been demonstrated in multiple mouse models of inflammatory, neuropathic, and visceral pain[6; 13; 16; 20], with no analgesic effect in mGlu5 knockout mice[20]. Additionally, two-week daily dosing did not result in tolerance to fenobam analgesia in mice[21]. Pre-clinical studies demonstrating the analgesic potential of fenobam, and the safety profile of fenobam observed in small clinical studies [2; 3; 12; 15; 22; 23; 31] encouraged us to utilize fenobam to test the hypothesis that mGlu5 modulates nociceptive sensitization in humans.

In prior human studies, fenobam was administered to only a limited number of subjects, and the disposition of this specific formulation was not thoroughly investigated. We sought to evaluate the pharmacokinetics of a single dose of fenobam, and to then test the analgesic effects of fenobam in a human experimental pain model. The first study was a dose escalation protocol in adult healthy volunteers. The highest dose administered (150 mg) was the highest single dose reportedly well tolerated in previous studies [2; 12; 23]. The second study evaluated 150 mg oral fenobam analgesia in the validated heath/capsaicin model of cutaneous sensitization [8; 24]. Capsaicin is an effective and selective activator of transient receptor potential vanilloid 1 (TRPV1), therefore we chose this model based on the suggested role of mGlu5 in modulating TRPV1-mediated sensitization [14]. Additionally, capsaicin is a commonly used model for afferent-induced secondary hyperalgesia in humans [27], thus providing a direct translation of our prior studies demonstrating the effect of fenobam on the second phase of the formalin test in mice [20; 21]. Our hypothesis was that the area of cutaneous sensitization[24] that would develop under the effect of fenobam would be reduced in size compared to the baseline area (measured on the training day) across time-points, while, with placebo, the area would not differ from baseline. Therefore, we expected the areas of sensitization to be significantly different under the two conditions (fenobam/placebo).

Given the reported anxiolytic properties of fenobam [12; 15; 22; 23; 25], we considered the possible implications of anxiolysis on pain modulation, and evaluated subject mood and affect. Moreover, because pre-clinical studies reported effects of mGlu5 modulation on working memory and cognitive performance[1; 13; 19; 26; 28], we also assessed memory and cognition before and after fenobam or placebo.

## Methods

All studies were carried out in accordance with ethical principles of Good Clinical Practice and the Declaration of Helsinki and its guidelines, and approved by Washington University Institutional Review Board, following submission of an Investigational New Drug (IND) application to the FDA (IND#117,989). Subjects provided written informed consent. The investigations were registered on ClinicalTrials.gov NCT01806415 and NCT01981395. Fenobam was manufactured under GMP guidelines by Scynexis, Inc. (Durham, NC) as powder and then compounded in gelatin capsules with lactose monohydrate at the Washington University in St. Louis investigational Pharmacy under strict adherence to USP 795.

### Study Setting

Both studies (“PK Study” and “Hyperalgesia Study”) were conducted in the Washington University Clinical Research Unit (CRU) and/or the Anesthesiology Human Studies Lab at Washington University. Participants in the two studies were screened and enrolled separately. None of the subjects participated in both of the studies here presented. Separate methods for each of the two studies follow here below.

## 1. PK Study. Pharmacokinetics and Side Effects of Oral Fenobam in Adult Healthy Volunteers

This was a randomized, double blind, single dose, parallel group, placebo controlled study to evaluate the pharmacokinetics and side effects of fenobam (Flow Chart – Fig.1A). The primary objective of this study was to obtain pharmacokinetic data after oral administration of 50, 100 and 150 mg of fenobam in groups of healthy individuals. A secondary objective was to compare the side effects of a single dose of 50 mg, 100 mg or 150 mg of fenobam to placebo.

**Fig. 1.**
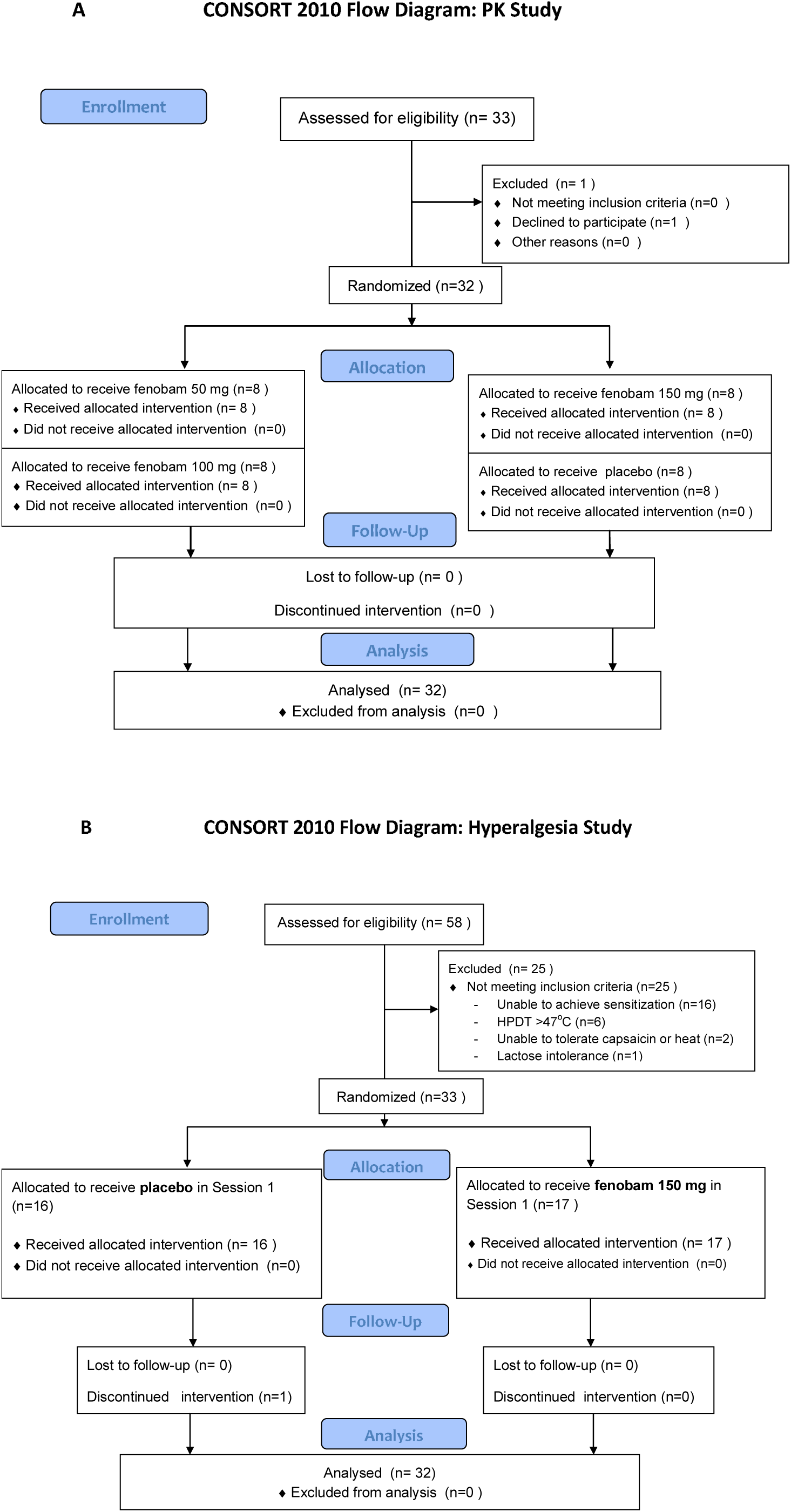
CONSORT Flow Diagrams: **A**. Study of the pharmacokinetics of fenobam (“PK study”); **B.** Study of the analgesic effects of fenobam on the heat/capsaicin-induced cutaneous hyperalgesia in adult healthy volunteers (“Hyperalgesia study”). The flow diagram of the “Hyperalgesia study” only shows “Session 1” of the originally intended “placebo-controlled crossover design” as the design was revised, and Session 2 was eliminated from analysis, following the identification of a significant carryover effect in the second study Session.

### Screening Session/Pre-study Period for the PK study

Potential candidates were screened by the PI (LFC) and by the Senior Research Coordinator (KF) according to the inclusion and exclusion criteria. Inclusion criteria were age 18-50 yr, good general health with no remarkable medical conditions (e.g., history of liver, kidney, heart, or lung disease), and BMI < 33. Exclusion criteria were medication use (prescription or non-prescription medications, vitamins, herbals, dietary and mineral supplements and grapefruit products during or within 14 days prior to study participation; excluding contraceptives), history of addiction to drugs or alcohol (prior or present addiction or treatment for addiction), pregnancy or nursing, history of lactose intolerance, and smoking. Study participants were asked to abstain from drinking alcohol for 24 hours before the study, and to abstain from eating and drinking after midnight on the night before the study. An initial visit included patient health self-assessment, medical history and physical examination. Vital signs were then recorded (heart rate, respiratory rate, blood pressure, temperature and oxygen saturation). Each subject who qualified for entry into the study on the basis of inclusion/exclusion criteria, agreement with informed consent and pre-study evaluation was assigned the next available patient number. Thirty-two subjects were enrolled, with 8 subjects assigned to each group. In this Phase I, single oral dose pharmacokinetic and side effects study, a formal sample size calculation was not performed and sample size justification was based on prior experience with clinical pharmacokinetic investigations.

### Randomization for the PK study

After successful completion of the screening session, a computer generated randomization schedule prepared by the Washington University in St. Louis investigational Pharmacy assigned subjects to administration of either 50 mg, 100 mg, 150 mg of fenobam ([1-(3-chlorophenyl)-3-(1-methyl-4-oxo-2-imidazolidinylidine) urea hydrate]) or placebo (lactose monohydrate 150 mg in gelatin capsules). Therefore each subject received a single dose of fenobam or placebo. Subjects and study personnel were blinded to drug/placebo by administration in identical gelatin capsules provided by the Investigational Pharmacy.

### Study Period for the PK study

Vital signs were recorded and a peripheral IV catheter was inserted in an arm for blood sampling. A baseline blood sample was used for a complete blood cell count and comprehensive metabolic panel. Fenobam or placebo was administered with a sip of water and venous blood samples were drawn before and after fenobam/placebo administration (at 0.5,1,2,3,4,5,6,10 and 24 hours). Blood samples were centrifuged at 3000 rpm for 10 minutes at room temperature. Plasma was then removed and frozen at – 20°C for later analysis. Vital signs were simultaneously recorded and subjects were queried for side effects (confusion, visual changes/blurred vision, dizziness, light-headedness, weakness, speech difficulties, abnormal cutaneous sensations, tingling, or numbness, nausea/vomiting, headache, metallic taste, hot flashes and any abnormal feelings). An additional blood sample was obtained on day 2 for a complete blood cell count and comprehensive metabolic panel.

One week after the study completion subjects were asked again to answer a questionnaire regarding any abnormal/unusual feeling they might be experiencing. The 1 week interview was conducted by phone.

### Laboratory Analysis for PK study

Plasma fenobam was quantified using tandem mass spectrometry, as previously described by our laboratory[20]. The calibration range was 6 to 16,000 ng/ml.

### Data Analysis for PK study

Descriptive statistics were generated for reported measures, and difference in distribution of baseline characteristics among groups administered fenobam 50mg, 100mg, 150mg or placebo were analyzed with the Kruskal Wallis H test using SPSS statistical software. The correlation between fenobam dose and number of side effects was investigated with Kendall’s tau coefficient of rank correlation (tau_b). Proportions of patients with side effects in the different groups were compared using Fisher’s exact test. For all analyses, careful attention was given to whether the data satisfied the distributional and model-specific assumptions of the procedures used.

C_max_, t_max_ and AUC_0-t_ were determined based on the measurements collected and using the statistical software package Sigma Plot 13.0 (Systat Software, Inc., San Jose, USA); Spearman’s correlation coefficient (r_s_) was calculated to measure the association between dose of fenobam administered and C_max_ and AUC_0-t_.

## 2. Hyperalgesia Study: Anti-Hyperalgesic Effect of a Single Dose of Fenobam on Heat/Capsaicin-Induced Cutaneous Hyperalgesia in Adult Healthy Volunteers

This study was designed as a randomized, double-blinded, placebo-controlled, two-way cross-over trial with 32 healthy volunteers who received either 150 mg fenobam or placebo (lactose monohydrate) and were then tested for cutaneous hypersensitivity using the heat/capsaicin model of cutaneous sensitization (Flow chart – Fig 1.B).

The primary outcome measure was suppression of the development of cutaneous hyperalgesia and allodynia around the area treated with heat/capsaicin. Heat/capsaicin application was timed for maximal sensitization to occur at the time of fenobam C_max_, as determined in the first investigation. Secondary outcomes evaluated were: Heat Pain Detection Thresholds (HPDT); pain scores after thermal stimulation of untreated skin, and side effects.

The study consisted of an initial screening/training session to evaluate subjects, establish eligibility and explain the study procedures, and two subsequent study sessions (placebo or fenobam, randomized order) that were conducted one week apart. During the two study sessions, blood samples were collected hourly for 7 hours after administration of fenobam, and fenobam plasma concentrations were then determined by tandem mass spectrometry. Measures of hyperalgesia and hypersensitivity were taken at regular time points during each study session, and included the area of cutaneous sensitivity to calibrated von Frey filament and to brush stroke stimulation, and Heat Pain Detection Thresholds (HPDTs). Assessments of mood/affect and cognitive function also occurred at the same time points. A short version [18] of the brief Positive and Negative Affect Scale (PANAS)[31] and Brief State Anxiety Measure (BSAM)[2] were utilized to evaluate mood and affect; the Letter and Number Sequencing Assessment (LNS), a subtest of the Wechsler Adult Intelligence Scale-Fourth Edition (WAIS-IV)[32] was used to evaluate working memory and cognitive function before and after fenobam or placebo. A complete overview of the study timeline is shown in Fig 2.

**Fig. 2.**
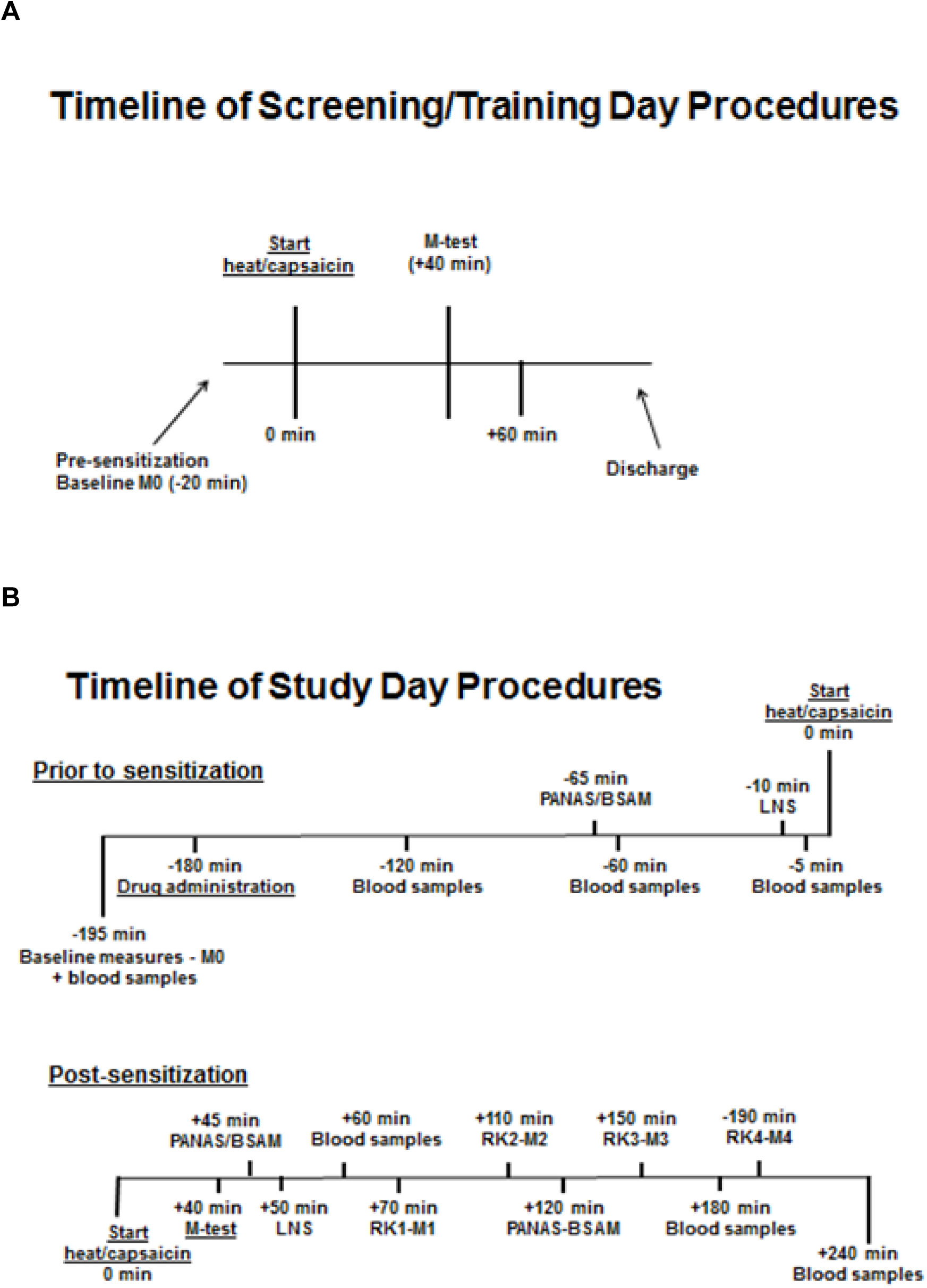
Timeline of the procedures on Screening/Trinaing Day (A) and Study Day (B). In each timeline, time points (min) are in relation to the “time 0” of start of the heat/capsaicin sensitization procedure; negative time points mark the pre-sensitization procedures and positive time points delineate the post-sensitization phase. **(A)** M = measurements; vF = von Frey; HPDTs= heat pain detection thresholds; PTS = pain at thermal stimulation. **(B)** PANAS = positive and negative affect scale; BSAM = Brief State Anxiety Measure; LNS = Letter and Number Sequencing; RK =rekindling of sensitization.

### Screening Session/Pre-study Period for the Hyperalgesia Study

Subjects who were potential candidates for the study were evaluated for study eligibility, according to the inclusion and exclusion criteria. These were the same as for the pharmacokinetic study, with the additional exclusions of anatomical malformation of an upper extremity, recent trauma or chronic lesions on either forearm, history of allergy or intolerance to capsaicin, and history of multiple drug allergies. An initial visit included a health self-assessment form, collection of medical history and physical examination. Vital signs were then recorded (heart rate, respiratory rate, blood pressure, temperature and oxygen saturation). Each subject who qualified for entry into the study on the basis of inclusion/exclusion criteria, agreement with informed consent and pre-study evaluation was assigned the next available patient number.

### Study Period for the Hyperalgesia Study

Subjects participated in three sessions: one screening/training session and two study sessions, each scheduled approximately one week apart. The *training session* was held on the same day as the above mentioned screening session. In this session the enrolled subjects were given a demonstration of the actual study procedures to be conducted during the study sessions (see below). The duration of the subjects’ participation was approximately four weeks, with the training session on day 1, the first study session within 14 days of the training session, and Session #2 approximately one week following Session #1. During the training session, subjects experienced the heat/capsaicin sensitization procedure and measurements of cutaneous sensitization as described in detail below. Subjects were withdrawn from the study if their heat pain detection threshold was greater than 47°C at baseline or they failed to develop an area of measurable cutaneous sensitization following heat and capsaicin stimulation. The training session was intended to familiarize subjects with the experimental procedures so that communication between experimenter and subject about the subjective experiences of painful stimulation could be as precise as possible. No drug was administered during the training session.

During the study sessions subjects underwent a complete set of sensitization procedures; mapping of the area of sensitization; measurement of pain from thermal stimulation and heat pain detection thresholds at baseline, and then multiple times following rekindling of the sensitization via heat application.

### Sample Size Estimates for the Hyperalgesia Study

The primary outcome measure was the size of the area of hyperalgesia and allodynia around the area sensitized by heat/capsaicin as quantified by cutaneous stimulation with foam brush strokes and a von Frey filament requiring 26 g of bending force. Preliminary data on the development of cutaneous sensitization with the heat/capsaicin model obtained from our group[4] enabled us to estimate the variability of the primary outcome measure, so that we could calculate the sample size sufficient to detect a 20% difference in the area of hyperalgesia in the two treatment conditions.

In the population of 15 subjects in the study referenced above[4], reproducing areas of skin sensitization to von Frey filament stimulation with the heat/capsaicin model in the same subjects 1 week apart, the within-day standard deviation of the area ranged from 26 (within day standard deviation of initial and final areas for the same subject on Session 2) to 30 (within day standard deviation of initial and final areas for the same subject on Session 1). The standard deviation of the difference between the areas for the same subject between days (Session 1 vs Session 2) was 13.

Based on these standard deviations, we calculated the sample size needed to detect a 20% difference in the area of sensitization measured in the two treatment conditions (80% power, alpha = 0.05, two-sided). A total of 32 subjects were estimated to be sufficient for this two-treatment crossover study. To detect a 30% difference in size of area of hyperalgesia between treatments, 15 subjects would be sufficient.

### Randomization for the Hyperalgesia Study

A computer-generated randomization schedule prepared by the Washington University in St. Louis investigational Pharmacy assigned subjects to fenobam 150 mg or placebo administration, and they were then crossed over to the other drug for the second session one week later.

### Study Procedures and Measurements for the Hyperalgesia Study

#### Administration Protocol

The heat/capsaicin hyperalgesia model combines heat stimulation (heat ramps from 32°C to 45°C at a rate of 1°C per second and hold at 45°C for 5 min) applied to a 9 cm^2^ area of skin on the forearm followed by topical low dose capsaicin (0.1% Capzacin-HP Cream) applied to the same area in such a way that the entire heated area was completely covered by a thick layer of cream, (approximately ¼ inch). The sensitization is then rekindled with subsequent applications of heat (40° for 5 min) at 35-45 minutes intervals. This procedure generates temporary pain and the sensory changes associated with peripheral and central sensitization for up to 4 hours[24]. Thermal stimulations were applied in a precise and controlled manner using a Medoc Advanced Thermal Stimulator (Medoc, Israel and North Carolina, USA) driving a 9 cm^2^ thermode. The thermode was a computer-controlled Peltier device that warms the skin from 32°C to a safety cutoff of 52°C in 1 C/sec increments.

#### Measurement of Pain at Thermal Stimulation, Pain Thresholds and Areas of Hypersensitivity

The following methods were used to induce and quantify pain and sensitization: 1) the pain intensity and area (cm^2^) of secondary hyperalgesia and allodynia induced by the heat/capsaicin model 2) heat pain detection threshold; 3) pain intensity produced by 1 minute 45°C thermal stimulation.

##### 1) Heat/Capsaicin Sensitization Procedure and Mapping an Area of Secondary Hypersensitivity

Sensitization was established by heating a 9 cm^2^ surface of the dominant forearm, and then applying approximately 14 ml of 0.1% capsaicin cream as described above. Subjects were asked to rate their pain on a Visual Analog Scale (VAS) at the start of the 30 min period and then for every 5 min until the cream was wiped off. At the end of the 30 min capsaicin application, measurements were performed to determine the areas of hypersensitivity and allodynia on the forearm. The borders of secondary mechanical allodynia and hyperalgesia were mapped using a 1 inch foam brush and a von Frey filament (26g bending force). Subjects were asked to close their eyes during these procedures. First, the brush was applied along four linear paths between the thermode outline and 1) the antecubital fossa, 2) the wrist joint, 3) the lateral aspect of the forearm in anatomical position, and 4) the medial aspect of the forearm. Stimulation started distant from the heated area and worked closer in 5.0 mm steps at 1sec intervals. Subjects were asked to say when the stimulation first became painful, and that location was marked. This procedure was then repeated with the von Frey filament. The area of hypersensitivity was calculated as the distance between the farthest points on the rostral/caudal axis multiplied by the distance between the farthest points on the medial/lateral axis (in cm^2^), then subtracting the heated area of the thermode (Fig. 3).

##### 2) Heat Pain Detection Threshold (HPDT)

Thresholds for heat pain detection were determined by using a thermal ramp protocol on a marked location on the volar surface of the forearm. The temperature applied through the Medoc thermode was increased from 32°C to the 52°C safety cutoff at 1°C/s. Subjects were requested to turn off the heated thermode by pressing a button at “the lowest temperature that they perceive as painful.” Four thermal ramps were performed 10 seconds apart; the values from the first ramp were excluded as they were consistently found to be lower than the following ramps, and the values from the remaining three ramps were averaged for each subject. To avoid testing individuals whose pain threshold approached the safety cutoff, subjects with HPDTs greater than 47°C were excluded from the study.

**Fig. 3.**
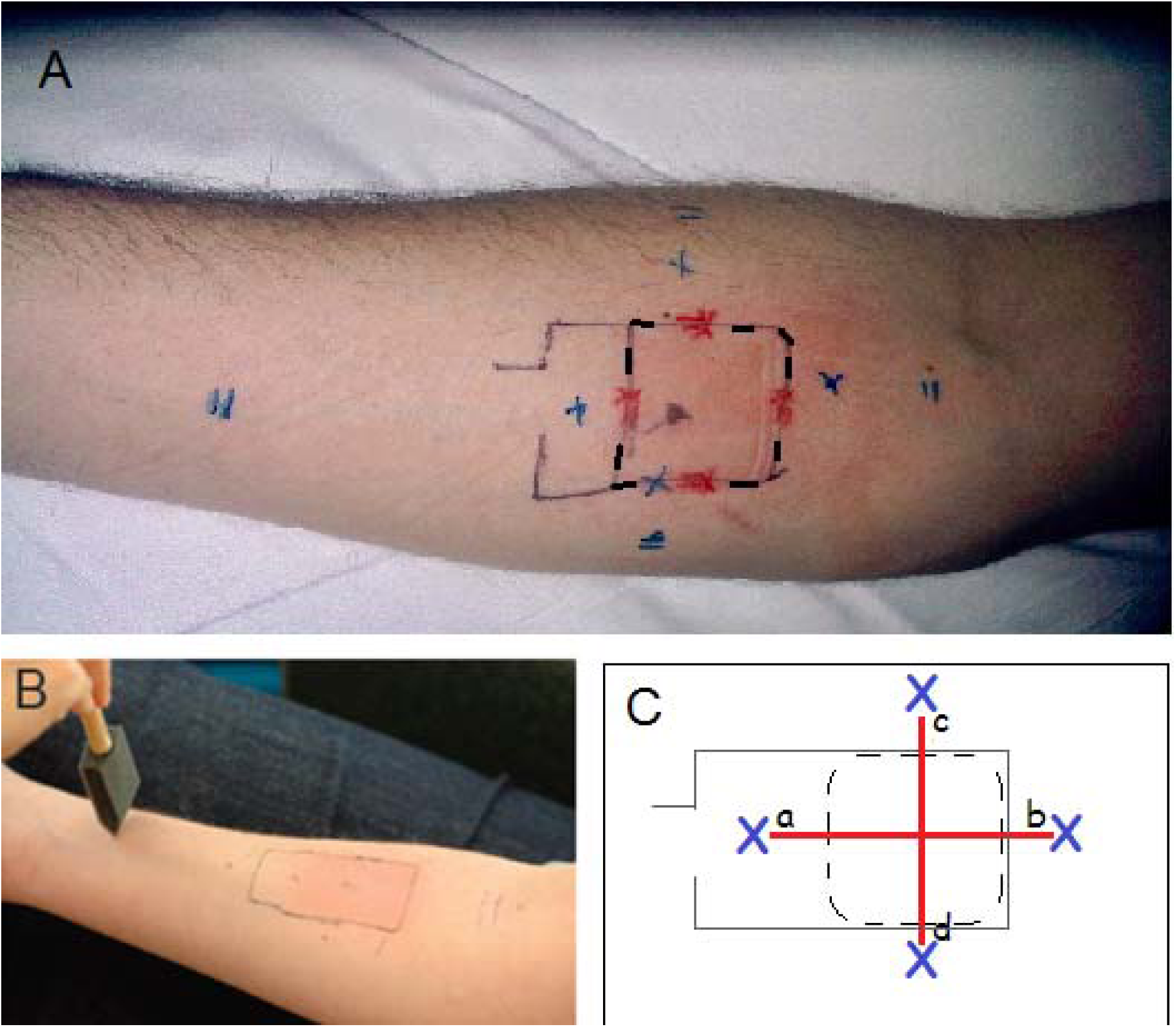
Measurement of cutaneous hypersensitivity after heat and capsaicin sensitization. **(A)** Thermode outline on a subject’s forearm and markings indicating points of change in sensation on the skin from “non-painful” to “painful” at foam brush (X) and von Frey filament (II) stimulation. The square marked with a broken line (- -) inside the thermode’s outline represents the heated surface of the thermode which measured 9 cm2. **(B)** Stimulation with brush strokes is shown as performed along a linear path between the wrist joint and the thermode outline. **(C)** Measurement of an area of hypersensitivity (brush area shown here): [(ab) × (cd)] – 9 cm^2^.

##### 3) Pain During Thermal Stimulation (PTS)

Acute pain was induced by a 1 min 45°C heat stimulus on a marked location on the upper non-dominant arm (deltoid). Subjects were asked to rate their pain intensity during the 1 min heat stimulus continuously using an electronic Visual Analog Scale (VAS) ranging from 0 to 100 where 0 indicates “no pain sensation” and 100 indicates “the most intense pain imaginable.”

### Rekindling Procedures

On drug study days (Session1 and 2), hypersensitivity was maintained by rekindling the site of heat/capsaicin application. This was accomplished by re-stimulating the previously treated skin four times at approximately 40-45 min intervals, with the thermode increasing from 32°C to 40°C) at a rate of 1° per second and held at 40°C for 5 min. Subjects rated their pain on a continuous visual analogue scale (VAS) during rekindling.

### Measurement/Evaluation of Mood and Affect Changes

Using a combination of Brief Positive and Negative Affect Scale (PANAS) and Brief State Anxiety Measure (BSAM)[18] we aimed to quantify changes from baseline in the subjects’ mood and affect following administration of the drug and after sensitization.

### Assessment of Cognitive Function

Changes in working memory (attention, concentration and mental control) were assessed with the Letter-Number Sequencing (LNS) test [30]. Changes in performance from baseline following administration of the drug and after sensitization were recorded.

### Data analysis

Our primary outcome measure was the size of von Frey and brush areas of cutaneous sensitization to heat and capsaicin.

A mixed within–between subject linear model approach using SAS Proc Mixed procedure was used to analyze the data. The mixed model used restricted maximum likelihood estimation for linear models with degrees of freedom adjusted using Kenward-Roger procedure. This model allowed controlling for potential confounders. Type III tests of fixed effects were used to evaluate the main effects of treatment group, time and interaction of treatment group with time.

The possibility of a carryover effect was explored by comparing baseline values between the two treatment Sessions in the same group (Wilcoxon signed ranks test), as well as by testing the sequence of randomization in the mixed model analysis. Since we found that there was a significant carryover effect, we analyzed separately data from Session 1 of the study. Independent samples t-test or non-parametric equivalent Wilcoxon rank sum test were used to explore differences in distribution of continuous level characteristics and baseline measures between subjects randomized to fenobam or Placebo in Session 1. In essence, the mixed model approach allowed us to explore the difference in size of von Frey area of sensitization through different time points and compare area sizes between the two treatment groups. The same analytical approach was also used for brush area and HPDT measures.

## Results

### 1. Pharmacokinetic Study. Pharmacokinetics and Side Effects of the mGlu5 Negative Allosteric Modulator Fenobam in Adult Healthy Volunteers

A total of 32 adult healthy volunteers (Table 1) were randomized to four groups to receive oral administration of placebo or fenobam (50, 100 or 150 mg). Fenobam plasma concentration curves after 50,100, and 150 mg oral doses are shown in Fig.4. C_max_, t_max_ and AUC_0-t_ median, minimum and maximum values for the three groups receiving fenobam are summarized in Table 2.

**Fig. 4.**
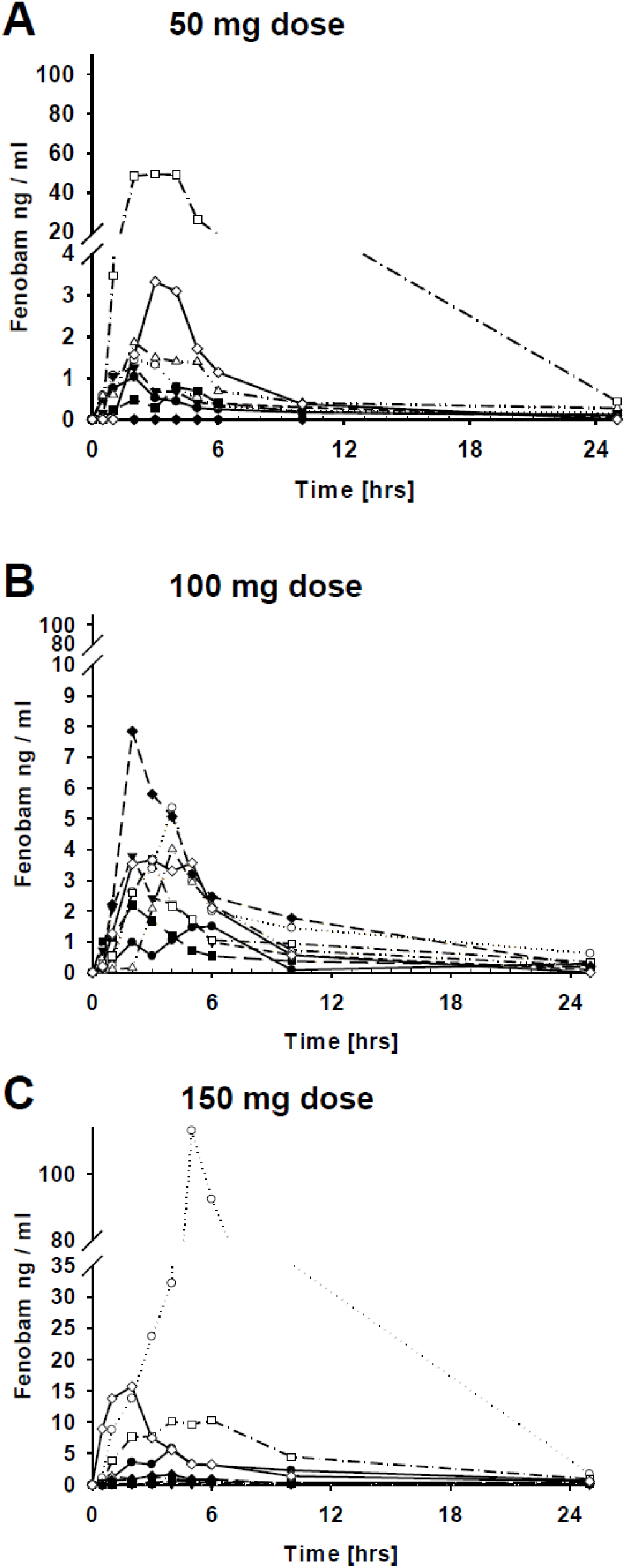
PK Study: Fenobam plasma concentration curves after 50 mg,100 mg, and 150 mg oral doses. Curves of individual subjects are shown in **A (50 mg), B (100 mg), and C (150 mg)**: each symobl/line combination represents a different subject for each group.

**Table 1.** PK Study: demographic information and baseline characteristics.

|  | All subjects<br>N=32 | Placebo<br>(n=8)<br>Median (Min-Max) | Fenobam<br>50 mg (n=8)<br>Median (Min-Max) | Fenobam<br>100 mg (n=8)<br>Median (Min-Max) | Fenobam<br>150 mg (n=8)<br>Median (Min-Max) |
| --- | --- | --- | --- | --- | --- |
| **Sex** |  |  |  |  |  |
| **Female** | 18 (56%) | 7 (87.5%) | 6 (75%) | 2 (25%) | 3 (37.5%) |
| **Male** | 14 (44%) | 1 (12.5%) | 2 (25%) | 6 (75%) | 5 (62.5%) |
| **Age** | 28 (20-49) | 29 (20-49) | 30 (21-35) | 24 (21-46) | 29 (21-40) |
| **Height (cm)** | 170 (157-198) | 170 (157-175) | 169 (163-180) | 176.5 (165-185) | 174 (160-198) |
| **Weight** | 77 (53-105) | 82 (66-95) | 71 (54-90) | 77 (54-98) | 78 (53-105) |
| **BMI** | 26 (19-33) | 28 (22-33) | 25 (20-29) | 25 (19-30) | 25 (20-30) |

**Table 2.** C_max_, t_max_ and AUC_0-t_ value medians (min; max) for the three groups of subject receiving 50 mg, 100 mg or 150 mg of fenobam.

| **Fenobam**<br>**Dose** | **C<sub>max</sub>**<br>**(ng/ml)** | **t<sub>max</sub>**<br>**(hrs)** | **AUC<sub>0-t</sub>**<br>**ng\*h/ml** |
| --- | --- | --- | --- |
| **50 mg** | **1.4 (0;48.4)** | **2 (2;4)** | **67.8 (0; 2405)** |
| **100 mg** | **2.9 (0.5;3.7)** | **3 (2;5)** | **167.4 (76.8;344.4)** |
| **150 mg** | **3.6 (0.1;32.2)** | **4 (2;6)** | **183.6 (10.8; 3862.8)** |

After oral administration of 50 mg fenobam, t_max_ was between 2 and 4 hours. C_max_ was extremely variable, between 0 and 48.4 ng/ml. T_max_ for the 100mg and 150mg oral doses was between 2 and 6 hours; for the 100 mg dose C_max_ were between 0.5 and 3.7 ng/ml, and for the 150 mg dose C_max_ were between 0.1 and 32.2 ng/ml. There was no significant association between fenobam dose administered and C_max_ (r_s_ = 0.236) or AUC_0-t_. (r_s_ = 0.295).

#### Safety and Tolerability

Fenobam was well tolerated up to the highest oral dose of 150 mg. Adverse events included headache, nausea, metallic or weird taste, and fatigue. All adverse events were described as “mild” by the subjects. Adverse events associated with either fenobam or placebo administration are presented in Table 3. No serious adverse events occurred in any subject. We found a weak negative correlation between fenobam dose and presence of side effects (Kendall’s tau_b −0.48; p=0.05) and the number of subjects with side effects was not significantly different between fenobam and placebo (Fisher Exact test, p= 0.82).

**Table 3.** Side Effects in subjects receiving placebo or different doses of fenobam. Rate of side effects in the groups **“Placebo”** and **“Fenobam 150 mg”** refer to subjects from both the PK study (8 subjects in each group) and hyperalgesia study (32 subjects receiving fenobam or placebo).

| **Reported Side Effect** | **Placebo**<br>**(n=40)**<br>**(%)** | **Fenobam 50 mg**<br>**(n=8)**<br>**(%)** | **Fenobam 100 mg**<br>**(n=8)**<br>**(%)** | **Fenobam 150 mg**<br>**(n=40)**<br>**(%)** |
| --- | --- | --- | --- | --- |
| **Headache** | **4 (10)** | **4 (50)** | **2 (25)** | **4 (10)** |
| **Nausea** | **3 (7.5)** | **1 (12.5)** | **1 (12.5)** | **0** |
| **Metallic taste /**<br>**dysgeusia** | **1 (2.5)** | **1 (12.5)** | **1 (12.5)** | **0** |
| **Fatigue** | **1 (2.5)** | **1 (12.5)** | **0** | **0** |
| **Drowsiness** | **11 (27.5)** | **0** | **0** | **10 (25)** |
| **Lack of**<br>**Concentration** | **2 (5)** | **0** | **0** | **0** |
| **Tingling** | **0** | **0** | **0** | **1 (2.5)** |
| **Dry mouth** | **3 (7.5)** | **0** | **0** | **1 (2.5)** |

### 2. Hyperalgesia Study. Effects of 150 mg Orally Administered Fenobam on the Development of Cutaneous Sensitization in the Heat Capsaicin Test

Demographic information and baseline characteristics of subjects who received the heat/capsaicin sensitization procedures are presented in Table 4. Disposition of oral fenobam 150 mg in this cohort is shown in Fig 5.

**Table 4.** Hyperlagesia Study: demographic information and baseline characteristics of subjects receiving placebo or fenobam in Session 1. HPDT = Heat Pain Detection Threshold.

| **Baseline characteristics**<br>**(on training day)** | **Placebo (n=15)**<br>**Median (Min-Max)** | **Fenobam (n=17)**<br>**Median (Min-Max)** |
| --- | --- | --- |
| **Age (y)** | **28 (22-34)** | **29(19-45)** |
| **Height(cm)** | **173(157-191)** | **170(157-188)** |
| **Weight (kg)** | **71(59-105)** | **79(65-90)** |
| **BMI** | **26 (20-32)** | **27(21-33)** |
| **Coffee cups** | **2(1-4)** | **1(0.5-3)** |
| **Forearm circumf. (cm)** | **26(23-33)** | **26(23-30)** |
| **HPDT(°C)pre-sensitization** | **43(38-47)** | **43(37-46)** |
| **HPDT(°C)post-sensitization** | **37(35-38)** | **37(35-39)** |
| **Brush Area (cm<sup>2</sup>)** | **78(45-143)** | **73(37-100)** |
| **vF area (cm<sup>2</sup>)** | **116(76-206)** | **90(81-124)** |
|  | **n (%)** | **n (%)** |
| **Sex** |  |  |
| **Female** | **8(53%)** | **11(65%)** |
| **Male** | **7(47%)** | **6(35%)** |
| **Daily Caffeine intake** |  |  |
| **Yes** | **10(67%)** | **11(65%)** |
| **No** | **5(33%)** | **6(35%)** |
| **Alcohol in last 6 months** |  |  |
| **Yes** | **14(93%)** | **15(88%)** |
| **No** | **1(7%)** | **2(12%)** |
| **Dominant Arm** |  |  |
| **Left** | **4(27%)** | **2(12%)** |
| **Right** | **11(73%)** | **15(88%)** |

**Fig. 5.**
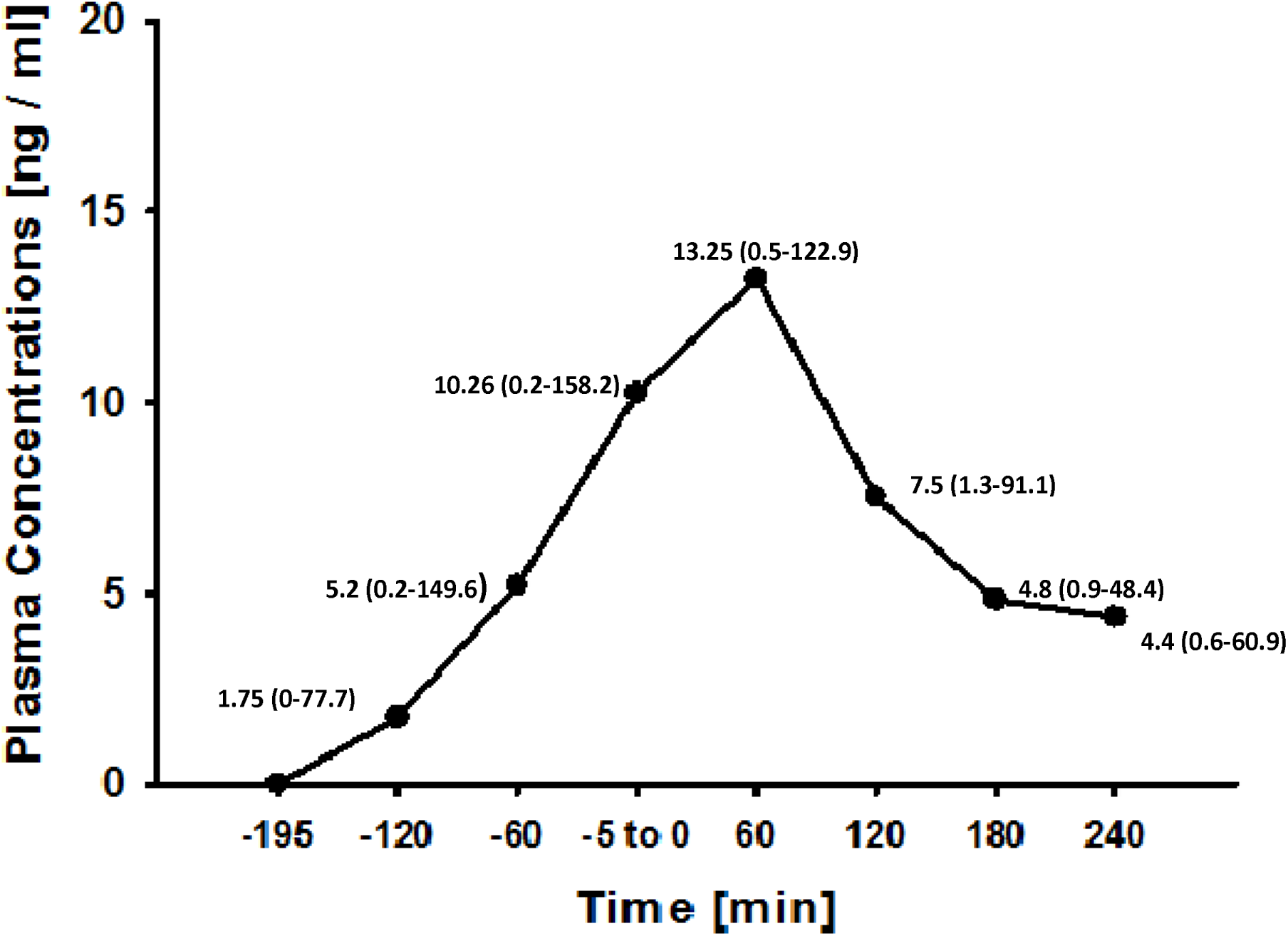
Hyperalgesia Study: fenobam plasma concentration following 150 mg oral dose. Median values (Min-Max) at different time points in relation to the start of the skin sensitization procedures (time 0) are shown.

#### Anti-hyperalgesic Effects

The primary outcome measure was size of areas of cutaneous sensitization to brush and von Frey filament. Taking into account the crossover design of the study, the presence of a carryover effect was explored by comparing baseline (M-test) area measures – i.e. immediately following sensitization with heat-capsaicin – between the 2 treatment sessions, as well as by testing the sequence of randomization in the mixed model analysis (SAS Proc Mixed procedure). We found a significant (p=0.001 for von Frey areas and p<0.001 for brush areas) and consistent (present in 26/32 subjects) reduction in von Frey and brush area measures at the first measurement in Session 2 compared to the same time-point (M-test) in Session 1, independent of treatment. Both von Frey and brush areas were reduced by 20% in Session 2 (fenobam) compared to Session 1 in the group that had received placebo in Session 1. In the group that had received fenobam in Session 1, von Frey areas in Session 2 (placebo) were reduced by 25%, and brush areas were reduced by approximately 50%. In light of these findings that we interpreted as the presence of a carryover effect of the pain testing paradigm, we focused our analysis on Session 1, with 17 subjects randomized to receive fenobam and 15 subjects randomized to receive placebo.

In Session 1, the overall (unadjusted) effect of fenobam on von Frey area compared to placebo was an average reduction in area size by 18.1 cm^2^ (95% C.I. −4.6 – 40.8). The overall effect on brush area was an average reduction by 13.9 cm^2^ (−2.4–30.3).

HPDTs (unadjusted results) were on average 0.4 °C higher in patients receiving fenobam compared to placebo (C.I. −0.8 to 1.6).

After controlling for area of sensitization to von Frey filament on the training day and alcohol use, which were found to be significant confounders of the drug effect, the change of von Frey area through different time points (M-test = Measurements taken immediately post sensitization, through M4= measurements taken after the 4th rekindling procedure; time-points illustrated in Fig.2) was significant, with the lowest values noted at time M4. These area reductions did not differ significantly between the placebo and fenobam groups. The average (all subjects) reduction of von Frey area at M4 compared to the area measured at M-test was 44.0 cm^2^ (95% CI: 28.2 to 59.8) in the placebo group, corresponding to 41% reduction from M-test, and 35.4 cm^2^ (95% CI: 20.4-50.4) in the fenobam group, corresponding to 35% reduction from M-test. Overall, subjects treated with fenobam had a von Frey area 0.76 cm^2^ smaller than subjects treated with placebo; however this difference was not statistically significant (95% CI:-16.03 to 17.56).

A similar pattern was also observed in brush area measurements. After controlling for area of sensitization on training day and forearm circumference (confounding variables of the drug effect in this model), the mean reduction of brush area at M4 compared to M-test was 46.9 cm2 (95% CI: 34.3 to 59.4) in the placebo group, corresponding to 70% reduction from M-test, and 44 cm^2^ (95% CI 32.0-55.9) in the fenobam group, corresponding to 71% reduction from M-test. Overall, subjects treated with fenobam had a brush area 7.5 cm^2^ smaller than placebo group with no statistical significance (95% CI:-7.1 to 22.1).

When von Frey and the brush areas measured immediately post-sensitization, at the single time point closest to the C_max_ of fenobam (= M-test on Study Day, Fig.2) were compared with areas obtained at M-test on the training day within the same group of subjects (but without fenobam or placebo administration), both areas were significantly reduced in size (median von Frey areas 74.6cm^2^ vs 90.1cm^2^ (17% reduction); brush areas 54.7cm^2^ vs 72.65 cm2; (25% reduction) p=0.025 and 0.028 respectively), while the areas measured at M-test of Session 1 in the placebo group were not statistically different from training day (median von Frey areas 112.4cm^2^ vs 115.8cm^2^ (3% reduction); brush areas 66.8 cm^2^ vs 77.9; (14% reduction) p = 0.069 and p = 0.1 (Fig.6A and 6B).

**Fig. 6.**
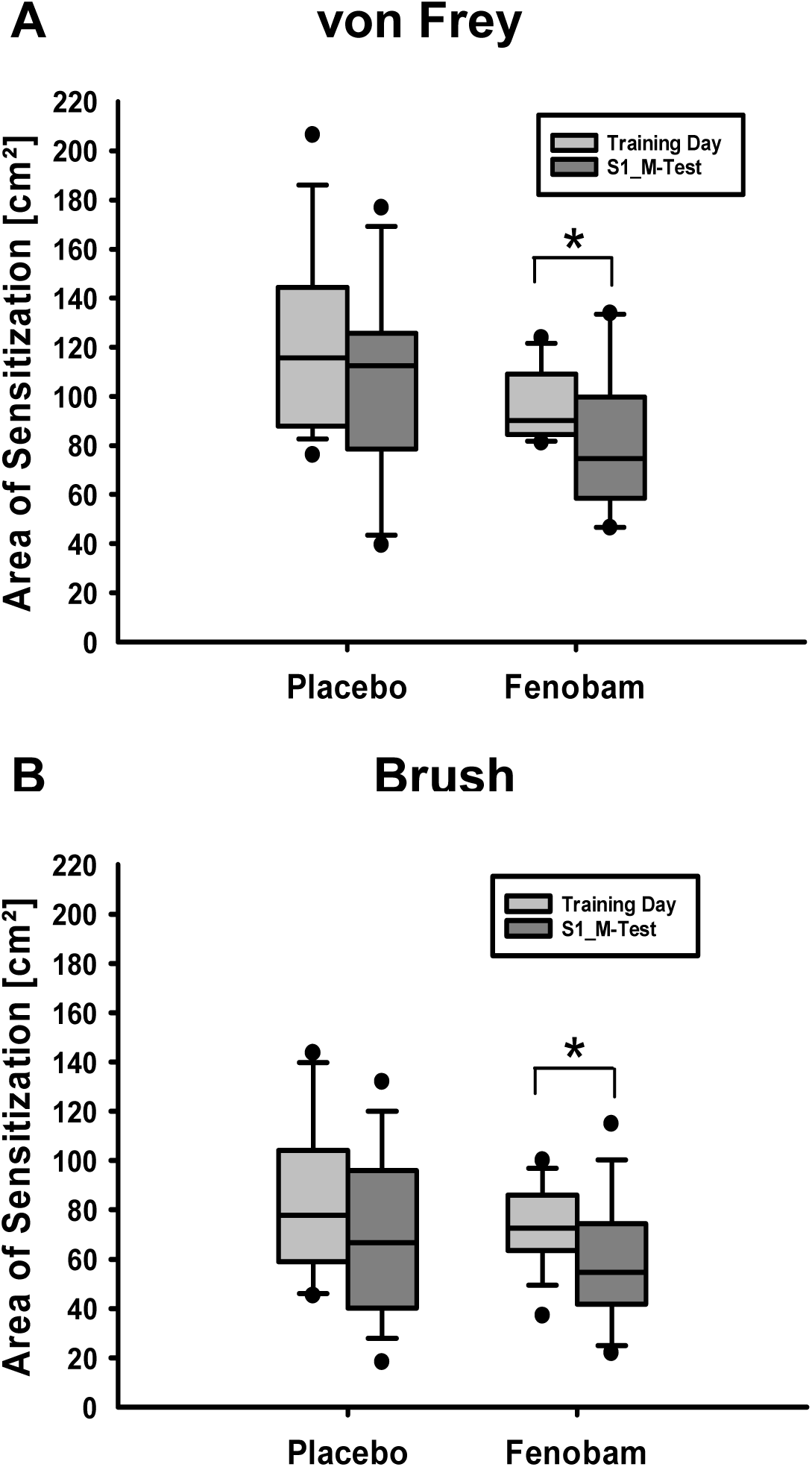
von Frey (A) and brush (B) areas of cutaneous hypersensitivity measured immediately after sensitization (M-test time-point) on Session 1 (S1) compared to areas on Training Day. Median; minimum; maximum values of areas; lower and upper quartiles are shown. The difference between median **Training Day** areas (no treatment) and median areas on **Session 1** is significant when fenobam is administered [*p = 0.025 (von Frey, panel A); *p = 0.028 (brush, panel B)] but not when placebo is administered.

After controlling for HPDT at baseline (Session 1), over all time points, mean HPDTs in the fenobam group were 0.2 °C higher than the placebo group; however this difference was not significant (95% CI −0.7 to 1.1)). At the M-test time-point there was no difference in HPDTs between the fenobam and placebo group.

#### Anti-nociceptive Effect

Overall after controlling for PTS at S1 baseline, there was no significant difference in maximum VAS (PTS MAX) scores recorded between the 2 groups and across all time-points (data not shown).

#### PNAS/BSAM and LNS

In a multi-level model to determine whether the mood of the subjects and their anxiety level varied with fenobam plasma concentration over time we could not observe any statistically significant effect of fenobam compared to placebo (not reported). Measures of cognitive function (attention, concentration and mental control) were also not significantly different between groups (not shown here).

## Discussion

Our study of the pharmacokinetic properties and side effect profile of fenobam after oral administration in human subjects showed highly variable disposition and t_max_ between 2 and 6 hours with marked inter-individual differences. These observations are similar to prior findings on the variability of fenobam disposition in a pilot study on a limited number of subjects with Fragile-X syndrome (FXS), wherein Berry-Kravis et al.[2] found that fenobam plasma concentrations were dose-dependent but variable, with mean C_max_ of 39.7 ±18.4 ng/ml at 180 min following oral administration of fenobam150 mg. In the Berry-Kravis study, three healthy volunteers (control group) had a mean peak plasma concentration of fenobam that did not differ from that seen in the FXS patients. The timing to mean peak plasma concentrations was similar in FXS patients and in healthy controls. Neither fenobam dose nor concentrations after 60 or 120 min correlated with improvement of the clinical outcome of interest (behavioral response).

In contrast to results obtained in pre-clinical studies by our group [20], which showed robust analgesic effects of fenobam in mice, we observed a transient reduction in the area of hypersensitivity at the time of peak fenobam plasma concentration, but did not observe any persistent anti-hyperalgesic or anti-nociceptive effect of fenobam compared to placebo in this cutaneous sensitization experimental pain model. The relatively unimpressive effect of fenobam in this human study compared to the rodent studies may be due to multiple factors, including the unknown relationship between plasma concentration and receptor occupancy (and accordingly the potential for insufficient fenobam dosing in the present study) or, of course, simply reduced drug effect in humans relative to rodents.

We are not aware of other studies in human that evaluated the effect of an mGlu5 negative allosteric modulator as an analgesic. In preclinical animal studies, mGlu5 has been shown to modulate pain-related plasticity at multiple levels of the pain neuraxis, and mGlu5 antagonists have demonstrated analgesic properties in a broad range of pain models [6; 14; 16; 20]. In an attempt to reproduce in human the same conditions in which we had observed an analgesic effect of fenobam in rodents, we utilized an experimental model that aimed to be the direct translation of rodent experimental pain models that we had used [20; 21; 27]. The single dose of fenobam that we administered in our human study was approximately equivalent to the dose that we administered in mice with analgesic effect (allometric scaling suggests that 30 mg/kg in mice corresponds to 2.4 mg/kg in human, with a total dose of 150 mg being the calculated dose for an ideal body weight of 60 kg) (see “Guidance for Industry”: “Estimating the Maximum Safe Starting Dose in Initial Clinical Trials for Therapeutics in Adult Healthy Volunteers” Published by the U.S. Department of Health and Human Services; Food and Drug Administration Center for Drug Evaluation and Research (CDER); July 2005).

We observed a reduction of von Frey area through different time points (M-test = immediately post sensitization, through M4= after the 4th rekindling procedure), with the lowest values noted at time M4. These area reductions did not differ significantly between the placebo and fenobam groups. This finding was consistent with previously reported progressive fading of areas of hyperalgesia over time[4]. The presence of a strong carryover effect of the sensitization model from Session 1 to Session 2, and possibly unexplained interactions between the carryover effects of both the sensitization model and of the drug, prevented us from taking advantage of the original cross-over design.

Before starting the study, we were aware of potential issues with the heat/capsaicin model, its reproducibility and presence of a carryover effect causing areas of sensitization in Session 2 to be smaller than Session 1 in a crossover design[4]. However, based on our pre-clinical data, we had anticipated that the magnitude of the analgesic effect of fenobam would be sufficiently large to detect despite the carryover effect of the model. A complete unknown here is the relative receptor occupancy by fenobam in the rodent and human studies. Additionally, despite the purported[8; 9; 24] adequacy of the washout period for fenobam demonstrated by PK analysis, we cannot exclude the possibility that an effect of fenobam on cutaneous sensitization in Session 1 could have affected the sensitization process in Session 2, one week later, when the same subjects were receiving placebo. Our inability to exclude such an effect is based on the observation that areas in Sessions 2 have been consistently found to be smaller than Session 1. A possible explanation of this phenomenon is that cutaneous sensitization one week before may affect the development of a “re-sensitization” the following week. If this is true, any effect of fenobam on the sensitization process in Session 1 may have an indirect, unpredictable influence on the following week’s process. A parallel comparison of two groups receiving fenobam in the two consecutive sessions or placebo in both could have provided more information on the effect of the drug versus the effect of the repeated exposure to heat and capsaicin on skin sensitization. Unfortunately we are not able to administer repeated doses of fenobam to the same subjects due to regulatory constraints.

Once a strong carryover effect was detected, the decision was made to examine data from Session 1 separately, an approach that has precedent [7; 17]. However this approach decreases the statistical power of the study and has been deemed “at risk of bias”[7; 10; 11]. We recognize this as a significant limitation of our study, and we focused the presentation and interpretation of our findings on 95% confidence intervals.

With the limitations noted above, in the heat/capsaicin cutaneous sensitization model, contrary to our hypothesis that the size of the area of hypersensitivity would remain different in the two treatment groups across subsequent time-points as fenobam concentration was decreasing, we found no difference between fenobam and placebo in the overall analgesic effect. However, our study design also took into account the possibility that there might be only a narrow window around the time of C_max_ to detect an effect of fenobam in our constrained experimental conditions, and that this effect would not last throughout the sensitization procedures. To maximize the sensitivity of the clinical model, we performed the preliminary PK study and then planned the skin sensitization procedures so that initial sensitization would coincide approximately with peak plasma fenobam concentration.

Plasma fenobam analysis confirmed that initial heat and capsaicin sensitization did coincide with peak plasma fenobam concentrations. By analyzing sensitization data from the single time-point closest to C_max_ of fenobam (M-test) we found that both von Frey and brush areas were significantly reduced in size compared to areas obtained at baseline on the training day in the same subjects, when no drug was administered, while areas measured at M-test in the placebo group were not statistically different from training day. While this single time-point finding was observed both in the von Frey and the brush areas, possible explanation for the absence of a <u>persistent</u> anti-hyperalgesic effect of fenobam might be the rapid elimination of fenobam compared to the duration of cutaneous sensitization.

Availability of validated models to reliably create experimental hyperalgesia in healthy volunteers is limited, but – from a safety standpoint -we considered necessary to test fenobam in human healthy volunteers first, before conducting a prospective randomized clinical trial in patients affected by a clinical pain condition. With these considerations in mind, we relied on the prior extensive use of the model, and our direct knowledge and experience with the heat/capsaicin sensitization procedures. However, our limited and constrained experimental conditions might have impaired our ability to detect a persistent effect on the areas of sensitization over time.

Obviously, it is also possible that, despite the encouraging pre-clinical data, fenobam, and possibly mGlu5 antagonists in general, may not have robust analgesic effects in human. In our opinion, a clear answer on this issue requires additional studies where compounds with improved pharmacokinetic properties, and known receptor engagement, are utilized.

In conclusion, in our limited experimental conditions with human healthy volunteers, we did not observe any clinically or statistically significant persistent analgesic (anti-hyperalgesic) effect of fenobam compared to placebo over the study time course. Fenobam, administered orally to human healthy volunteers at a single dose of 50, 100 or 150 mg, showed highly variable disposition and t_max_ between 2 and 6 hours with marked inter-individual differences.

As part of our limitations, we found a significant carryover effect on the second study day, which led us to use a different data analysis strategy than was originally planned. Moreover, given the significant limitations introduced by the highly variable plasma exposure of fenobam, we feel that attempts to assess the potential utility of mGlu5 modulation for pain necessitate use of a compound with improved pharmacokinetics and known target engagement. Prospective randomized clinical trials with such an improved molecule are needed to clarify the role of mGlu5 modulation in the development and maintenance of acute and chronic pain conditions in human.

## Acknowledgements

The authors wish to thank Dr. Eric Lenze for his contribution as a consultant on the evaluation and measurement of mood and affect changes and measures of cognitive function and Alicia Neiner for the quantification of fenobam in plasma.

None of the authors has reported any conflict of interest, other than the financial support as reported below.

